# RAB6 and microtubules restrict secretion to focal adhesions

**DOI:** 10.1101/382176

**Authors:** L. Fourrière, A. Kasri, N. Gareil, S. Bardin, J. Boulanger, R. Sikora, G. Boncompain, S Miserey-Lenkei, F. Perez, B. Goud

**Affiliations:** Institut Curie, PSL Research University, CNRS, UMR 144, Dynamics of Intracellular Organization laboratory, 26 rue d’Ulm, 75248 Paris cedex 05, France; Institut Curie, PSL Research University, CNRS, UMR 144, Molecular Mechanisms of Intracellular Transport laboratory, 26 rue d’Ulm, 75248 Paris cedex 05, France; Institut Curie, PSL Research University, CNRS, UMR144, Cell and Tissue Imaging Facility (PICT-IBiSA), F-75005, Paris, France

**Author notes:** Equal contributions (L. Fourrière, A. Kasri, G. Boncompain, S Miserey-Lenkei, F. Perez, B. Goud).

## Abstract

To ensure their homeostasis and sustain differentiated functions, cells continuously transport diverse cargos to various cell compartments and in particular to the cell surface. Secreted proteins are transported along intracellular routes from the endoplasmic reticulum through the Golgi complex before reaching the plasma membrane along microtubule tracks. Using a synchronized secretion assay, we report here that exocytosis does not occur randomly at the cell surface but on localized hotspots juxtaposed to focal adhesions. Although microtubules are involved, the RAB6-dependent machinery plays an essential role. We observed that, irrespective of the transported cargos, most post-Golgi carriers are positive for RAB6 and that its inactivation leads to a broad reduction of protein secretion. RAB6 may thus be a general regulator of post-Golgi secretion.

## INTRODUCTION

To reach the cell surface, secreted proteins are transported along intracellular routes from the endoplasmic reticulum through the Golgi complex. Cargos exit the Golgi complex in transport carriers that use microtubules to be addressed rapidly to the plasma membrane before exocytosis. Transmembrane proteins are then exposed at the plasma membrane while soluble cargos are released in the extracellular space. Whether delivery of cargos occurs randomly or at specific sites of the plasma membrane is still unclear and the mechanisms that direct exocytosis are still unknown. Microtubules were described to be captured and stabilized by focal adhesions (Kumar et al., 2009). Their targeting to focal adhesions is driven by +TIPs (microtubule plus-end tracking proteins), such as APC, EB and CLASP, which ensure their physical contacts (Lansbergen et al., 2006; Akhmanova and Steinmetz, 2008; Kumar et al., 2012; Stehbens et al., 2014). Additionally, microtubules are linked to the actin network which is a structural component of focal adhesions (Palazzo and Gundersen, 2002). Notably, microtubules are involved in the regulation of the distribution and dynamics of adhesion sites (Small et al., 2002; Stehbens and Wittmann, 2012; Etienne-Manneville, 2013).

CLASPs (cytoplasmic linker-associated proteins) interact at the plasma membrane with a protein complex made of LL5β, a PI3P-binding protein, and ELKS (also known as RAB6IP2). ELKS is an effector of the Golgi-associated RAB6 GTPase (Monier et al., 2002), which regulates several anterograde and retrograde trafficking pathways to and from the Golgi complex, as well as Golgi homeostasis (Goud, 1999; White et al., 1999; Grigoriev et al., 2007; Mallard, 2002). In particular, RAB6 was shown to be involved in the targeting of post-Golgi vesicles containing the secretory markers VSV-G (vesicular stomatitis virus glycoprotein, a type I transmembrane protein), and NPY (neuropeptide Y, a soluble protein) to ELKS-enriched regions of the plasma membrane (Grigoriev et al., 2011; Miserey-Lenkei et al., 2010). RAB6 has been also shown to regulate the secretion of TNFα in macrophages (Micaroni et al., 2013), and the trafficking of herpes simplex virus 1 (HSV1) (Johns et al., 2014). However, thus far, no systematic study has been performed to characterize the cargos present in RAB6-positive vesicles.

The aim of this study was to investigate the spatial organization of post-Golgi trafficking of a variety of anterograde cargos in non-polarized cells. To this end, we combined the Retention Using Selective Hooks (RUSH) assay (Boncompain et al., 2012) to synchronize anterograde transport of cargos and the Specific Protein Immobilization (SPI) assay to map precisely the sites of arrival of the cargos at the plasma membrane. We show that cargos are transported along microtubules to hotspots of secretion, which are juxtaposed to focal adhesions. Moreover, we found that RAB6-dependent post-Golgi machinery plays a key role in this process and that RAB6 could be a general regulator of post-Golgi secretion.

## RESULTS

### Exocytosis takes place in restricted areas, close to the adhesion sites

Secretion of newly-synthesized proteins along the secretory pathway occurs continuously in cells. The RUSH system offers the possibility to synchronize the intracellular transport of cargos fused to the Streptavidin-binding-peptide (SBP) upon addition of biotin in the culture medium (Boncompain et al., 2012). With this system, it is possible to monitor a wave of secretion of a selected cargo and analyze its transport to the cell surface. Using the RUSH assay, we studied the synchronous secretion of diverse cargos: Collagen type X (ColX), VSV-G, secretory soluble EGFP (ssEGFP), gp135 (podocalyxin), CD59 and placenta alkaline phosphatase (PLAP), two GPI-anchored proteins, and tumor necrosis factor alpha (TNFα). Figure 1A illustrates RUSH-based transport monitoring using ColX as a cargo. As expected, before biotin addition, ColX was retained in the endoplasmic reticulum (ER) (Figure 1A, 0 min). Upon biotin addition, ColX left the endoplasmic reticulum, reached the Golgi apparatus within 10 min post-release and was then exocytosed at the plasma membrane. About 35 min after biotin addition, most of ColX had been secreted into the medium and almost no signal remained in cells. Time-lapse imaging and temporal projection after Golgi exit suggested that exocytosis did not occur randomly at the cell surface but in preferred domains (Figure 1A, Supp Movie 1). However, because ColX is a soluble secretory protein, a significant fraction of released proteins diffuses out, which may lead to underestimated levels of exocytosis at these preferred sites. To prevent its diffusion after release, we set-up an assay that we named Selective Protein Immobilization (SPI). In this assay, a GFP moiety is fused to soluble secretory factors or to the luminal part of membrane-bound cargos, and prior to seeding the cells, coverslips are coated with anti-GFP antibodies (Figure 1B). The interaction between the coated anti-GFP antibodies and the GFP moiety fused to the cargos reduces the diffusion speed of the cargos and eventually immobilizes them. This enables the local accumulation of secreted proteins that were released over an extended period of time. Combination of the RUSH and SPI assays thus provides a complete overview and localized history of the secretion of a selected cargo. Using SPI, and in contrast to Figure 1A, Figure 1C and Supp Movie 2 show a strong accumulation of secreted ColX visible 35 min after biotin addition. The presence of hotspots of ColX secretion confirmed that some domains of the plasma membrane seemed unable to support exocytosis while others were very active. The localization of the active domains was reminiscent of focal adhesion (FA) sites. We used cells expressing paxillin-mCherry, which localizes to FA (Turner, 1998; Turner et al., 1990), to monitor the synchronized transport of ColX combined with SPI and we found that secreted ColX was clearly enriched on FAs (Figure 1D). A similar result was obtained for another soluble cargo, ssEGFP, although it appeared more diffuse at the plasma membrane, likely due to rapid diffusion and/or less efficient capture by the antibody (Figure 1D).

**Figure 1:**
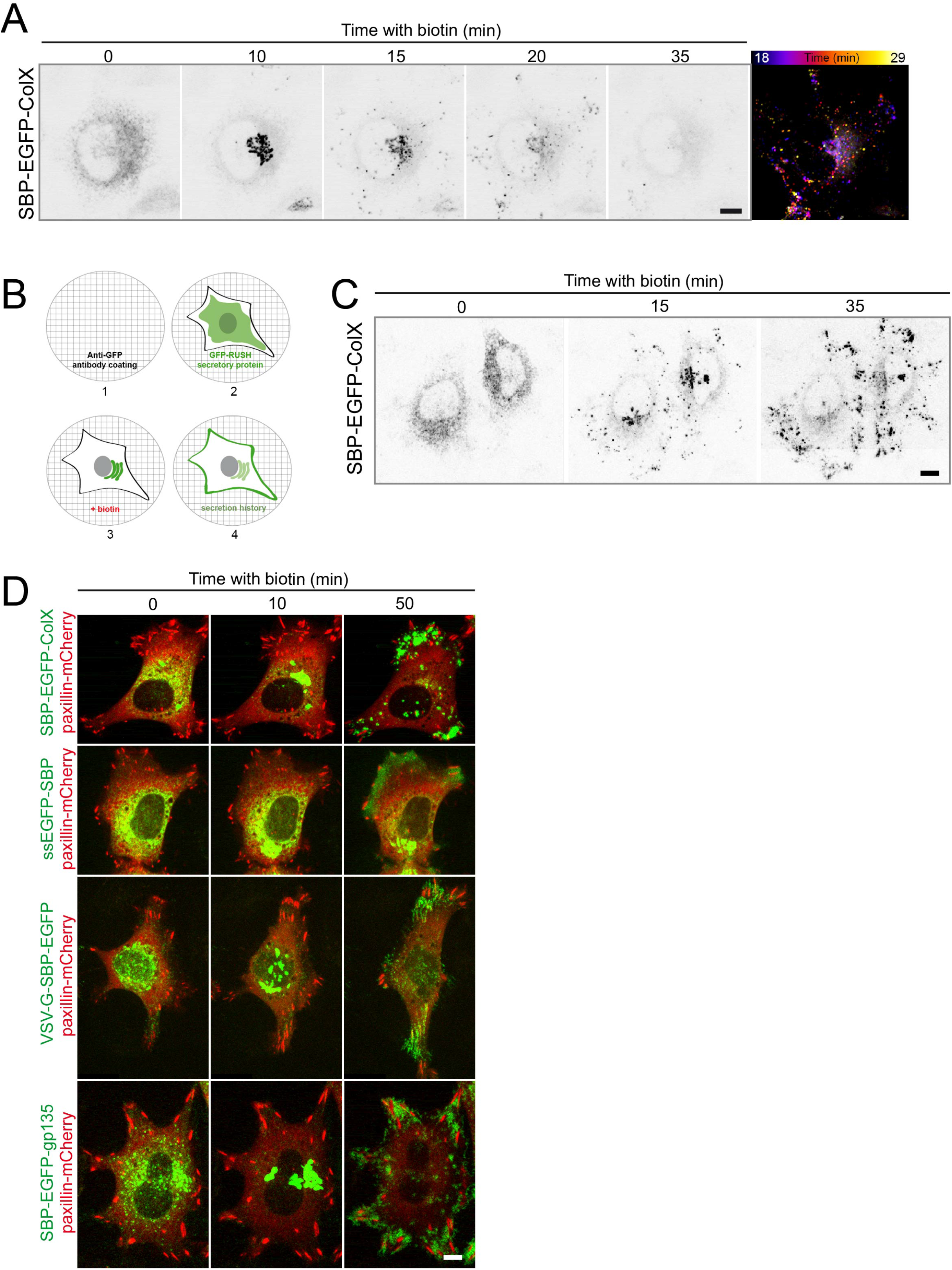
Local exocytosis close to adhesion sites of the cells. (**A**) HeLa cells stably expressing SBP-EGFP-CoIX were incubated with biotin for the indicated time (in min). Real time pictures were acquired using a spinning disk microscope and pictures were acquired at the indicated time. Temporal projection (right image) was performed for the SBP-EGFP-ColX signal between 18 and 29 min of trafficking. (**B**) Description of the SPI assay. 1. A coverslip is coated with an anti-GFP antibody. 2. The GFP-RUSH cell line is seeded on the coverslip. 3. Addition of biotin allows the trafficking of the GFP-RUSH-cargo. 4. Interactions between the anti-GFP and the GFP of the cargo (transmembrane or secreted) allow the capture of the cargo and provide a picture of the history of the secretion. (**C**) Trafficking of SBP-EGFP-ColX with an anti-GFP coating (SPI assay). HeLa cells stably expressing SBP-EGFP-ColX were incubated with biotin for the indicated time (in min). Real time images were acquired using a spinning disk microscope at the indicated time. (**D**) HeLa cells were transfected with SBP-EGFP-ColX, ssEGFP-SBP, VSV-G-SBP-EGFP or SBP-EGFP-gp135 and paxillin-mCherry. Coverslips were coated with an anti-GFP coating (SPI assay). Cells were observed by time-lapse imaging using a spinning disk microscope and pictures were acquired at the indicated time. Scale bars: 10 μm.

Similar experiments were performed with membrane-bound cargos like VSV-G, gp135, TNFα, and E-cadherin adapted to the RUSH assay. Although no particular enrichment was observed in normal conditions (Supplementary Figure 1), probably due to a rapid diffusion of secreted cargos in the plane of the plasma membrane, topologically-restricted secretion was observed using SPI. As for secreted cargos, exocytosis of VSV-G and gp135 also occurred on hotspots localized to FAs (Figure 1D). The same results were obtained when monitoring E-cadherin and TNFα secretion (data not shown).

The combination of the RUSH and SPI assays thus demonstrated the existence of secretion hotspots close to focal adhesions for soluble and membrane-bound proteins.

### Exocytosis is directed between focal adhesions

Next, real-time analysis of exocytic events was carried out using total internal reflection fluorescence microscopy (TIRF). In agreement with results obtained with the SPI set-up, we detected the frequent occurrence of exocytic events in certain regions of the plasma membrane, while other zones were seemingly silent (Figures 2A, B). Moreover, dual-color TIRF microscopy revealed that exocytosis did not exactly occur on focal adhesions, but juxtaposed to them, as confirmed by proximity measurement between the secretion puffs and paxillin signal (Figure 2C).

**Figure 2:**
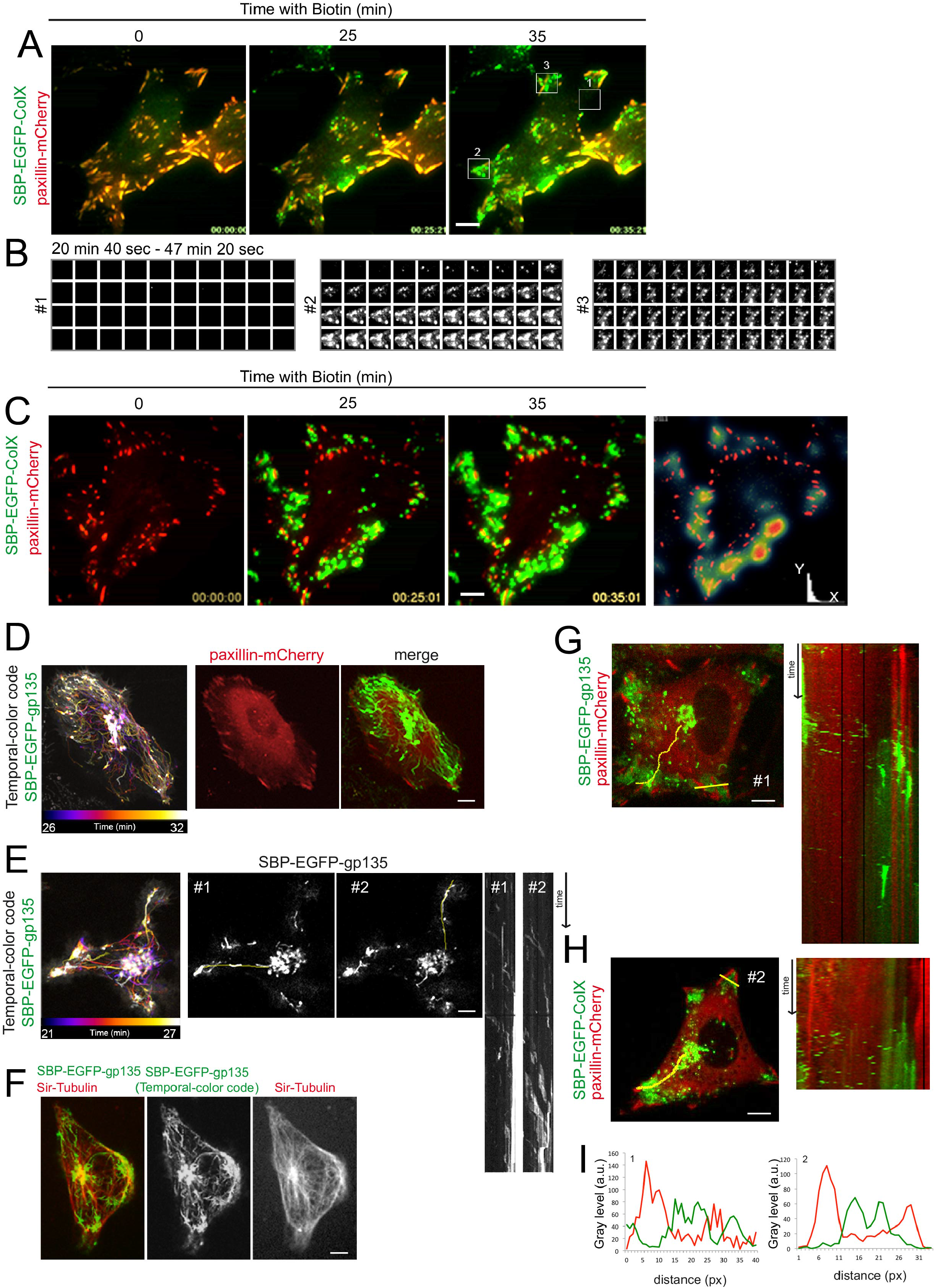
Exocytosis is directed between focal adhesions. (**A**) HeLa cells were transfected with SBP-EGFP-ColX and paxillin-mCherry. Real time pictures were acquired using a TIRF microscope and pictures were acquired at the indicated time. The TIRF angle was chosen based on the paxillin-mCherry signal. Biotin was added at time 0. (**B**) Areas 1, 2 and 3 are fixed images taken every 40 seconds between 20 min 40 s and 47 min 20 sec of ColX trafficking. Area 1 is a control condition without ColX signal at the plasma membrane. Areas 2 and 3 show puffs of secretion in close proximity to focal adhesions. (**C**) Coverslips were coated with an anti-GFP antibody (SPI assay). Real time images were acquired using a TIRF microscope and pictures were acquired at the indicated time. The TIRF angle was determined based on the paxillin-mCherry signal. Biotin was added at time 0. Right panel: Proximity map. Distance from the paxillin marker (x) was calculated at each position in function of the frequency of the secretion puffs of SBP-EGFP-ColX (y). (**D-H**) HeLa cells were transfected with SBP-EGFP-gp135 (E, F) or SBP-EGFP-gp135 and paxillin-mCherry (D, G) or SBP-EGFP-ColX and paxillin-mCherry (H). Coverslips were coated with an anti-GFP antibody (SPI assay). Microtubules were stained with SiR-tubulin (F). Cells were observed by time-lapse imaging using a spinning disk microscope and images were acquired at the indicated time. Biotin was added at time 0. After a 20 min incubation with biotin, SBP-EGFP-gp135 localizes in the Golgi apparatus and fast acquisition imaging was performed. Between 26 and 32 min (D) or 21 and 27 min (E), a temporal projection of EGFP-gp135 signal was performed using the Fiji software and is represented with a temporal-color code (left panel). Kymographs (time space plots) were performed on the indicated lines using the Fiji software. (**I**) Intensity profiles of SBP-EGFP-gp135 (G) or SBP-EGFP-ColX (H) (in green) and paxillin-mCherry (in red) at the plasma membrane were performed using the Fiji software at the indicated lines (#1 (G) and #2 (H)). px: pixels. Scale bars: 10 μm.

To explain how such a restriction of exocytosis may occur, we envisioned two non-exclusive hypotheses. On one hand, directed transport to focal adhesions may bias release toward adhesion domains. On the other hand, the factors essential to sustain exocytosis may only be present at the hotspots. To test the first hypothesis, we performed fast imaging (using a ~200 ms frame rate) of synchronized post-Golgi transport of gp135 to detect potential privileged tracks that may direct transport toward hotspots. Temporal projections revealed that gp135 *en route* from the Golgi complex to the cell surface used direct tracks toward focal adhesions (Figures 2D, E). Microtubules are involved in the regulation of the distribution and dynamics of adhesion sites, and can be captured and stabilized by focal adhesions (Small et al., 2002; Stehbens and Wittmann, 2012; Etienne-Manneville, 2013). Accordingly, we observed a co-occurrence of transport tracks with the microtubule network (Figure 2F), suggesting that a subset of microtubules targeting focal adhesions is used to release cargos. Kymographs drawn along the tracks (in yellow) indicated that the same tracks are used several times by distinct transport carriers coming from the Golgi apparatus in the direction of the plasma membrane (Figures 2G, H). While microtubules are abundant in cells, transport carriers thus seem to select microtubule subsets (Figures 2E, F). Kymographs and line scans also illustrated that secretion of cargo does not occur on focal adhesions but very close to them, in agreement with TIRF experiments (Figures 2G-I). This may be explained by steric hindrance due to the abundance and tight arrangement of proteins at focal adhesions and/or to specific localization of targeting/fusion factors.

Altogether, the above results indicate that secretory vesicles use preferential and direct microtubule-based routes for transport to secretion hotspots.

### ELKS and RAB6-dependent arrival of secreted cargos at exocytosis hotspots

Microtubules are attached to the cell cortex via the microtubule-stabilizing proteins CLASPs, which interact with a protein complex made of LL5β and ELKS (also known as RAB6IP2) (Lansbergen et al., 2006). We therefore investigated the presence of ELKS in secretion hotspots. As shown in Figure 3A, GFP-ELKS was enriched in zones of the plasma membrane close to FA where immobilized ColX is detected following 30-45 min biotin addition. In cells depleted for ELKS by siRNAs, the pattern of secreted proteins appeared more diffuse although secretion was still biased toward the regions near FA (Figures 3B, C). This suggests that ELKS, along with microtubules, contributes to the targeting the secretory vesicles to hotspots.

**Figure 3:**
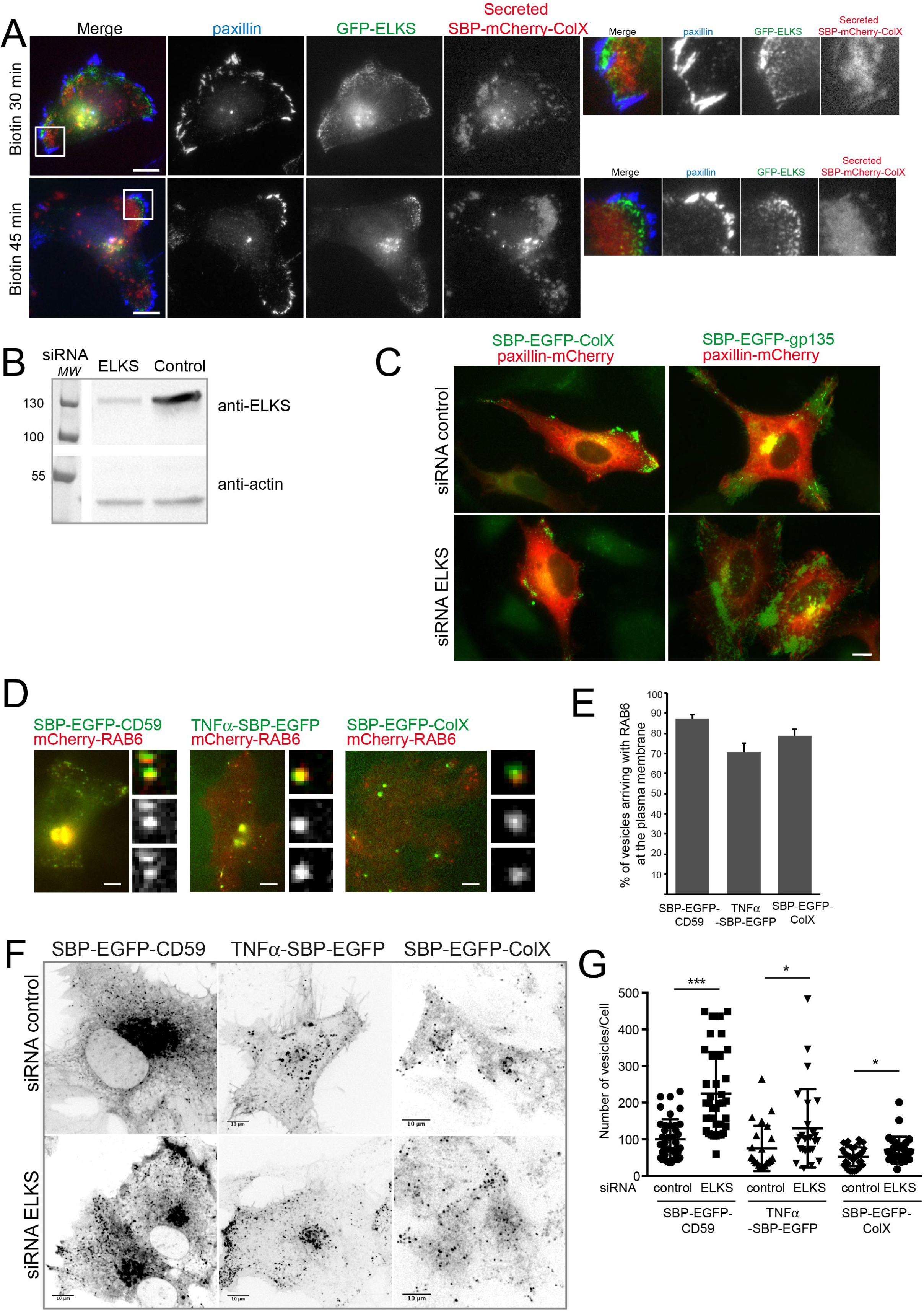
ELKS and RAB6-dependent arrival of secreted cargos at exocytosis hotspots. (**A**) HeLa cells were transfected with SBP-mCherry-ColX and GFP-ELKS. After 30 or 45 min incubation with biotin, cells were processed for immunoflorescence and stained with an anti-paxillin antibody. Coverslips were coated with an anti-GFP antibody (SPI assay). Higher magnifications of the images are shown on the right. (**B**) HeLa cells were treated with control or ELKS siRNA and processed for western-blotting. ELKS signal was revealed using an anti-ELKS antibody. Actin signal was used as a loading control. (**C**) HeLa cells were treated with control or ELKS siRNA and then co-transfected with SBP-EGFP-ColX or SBP-EGFP-gp135 and paxillin-mCherry. After a 45 min treatment with biotin, cells were fixed. Representative images are displayed. (**D**) HeLa cells co-expressing mcherry-RAB6 together with SBP-EGFP-CD59, TNFα-SBP-EGFP and SBP-EGFP-ColX were incubated for 30 min with biotin to allow cargos to leave the ER and reach the plasma membrane and then imaged using 3D-TIRF. Representative images taken from the movies are displayed. (**E**) Quantification of the percentage of RAB6-positive vesicles arriving at the plasma membrane and containing SBP-EGFP-CD59, TNFα-SBP-EGFP or SBP-EGFP-ColX. Cells were treated as indicated in D (mean ± SEM, n= 217-325 vesicles from 14-18 cells). (**F**) HeLa cells expressing SBP-EGFP-CD59, TNFα-SBP-EGFP and SBP-EGFP-ColX were treated for 3 days with control or ELKS siRNAs. Cells were incubated for 60-90 min with biotin to allow cargo release from the ER and its arrival to the plasma membrane. Representative images taken from movies. (**G**) The number of vesicles per cell were quantified using ImageJ (mean ± SEM, n= 23-39 cells). *** p< 10^−4^, * p< 0.05 (Student’s t-test). Cells were treated as indicated in F. Scale bars, 10 μm.

ELKS is a RAB6 effector (Monier et al., 2002), and was shown to be involved in the docking of RAB6-positive secretory vesicles containing VSV-G and neuropeptide Y (NPY) to the plasma membrane (Grigoriev et al., 2007). In addition, the very first study on the dynamics of GFP-RAB6 in living cells reported the presence of peripheral RAB6-positive structures near focal adhesions (White et al., 1999). We thus tested whether RAB6 was also associated with the transport carriers containing the cargos tested in this study, specifically ColX, TNFα and CD59. First, we used TIRF to investigate in cells co-expressing mCherry-RAB6 and either GFP-tagged ColX, TNFα or CD59, whether RAB6-positive vesicles arriving and fusing at the plasma membrane contained these cargos. We observed that about 80% of vesicles arriving at the plasma membrane and containing one of these cargos were positive for RAB6 (Figures 3D, E). In ELKS-depleted cells, in agreement with data reported previously (Grigoriev et al., 2007), we observed an accumulation of SBP-EGFP-CD59, TNFα-SBP-EGFP, or SBP-EGFP-ColX positive vesicles at the cell periphery as well as an increase in the total number of cytoplasmic vesicles (Figures 3F, G). Altogether, these data show that RAB6 and ELKS play a role in the docking and fusion of cargo-containing secretory vesicles with the plasma membrane.

### RAB6 associates with post-Golgi carriers containing GPI-APs, TNFα and ColX

We next investigated whether RAB6 was associated with post-Golgi carriers containing CD59, TNFα or ColX *en route* to the plasma membrane. Using live cell imaging, we performed a detailed analysis of the extent of co-localization between mCherry-RAB6 and several cargos during their transport (SBP-EGFP-CD59, TNFα-SBP-EGFP or SBP-EGFP-ColX). Following a 30 min incubation with biotin, we observed that 80% of vesicles containing one of the cargos were RAB6-positive (Figures 4A, C). By contrast, only 2-5% of RAB5-positive endosomal vesicles, used here as a negative control, were positive for CD59, ColX or TNFα in these conditions (Figures 4B, C). As expected, co-localization between RAB6 and cargos was minimal (10%) in post-ER compartments reached 10 min after biotin addition (Supplementary Figure 2).

**Figure 4:**
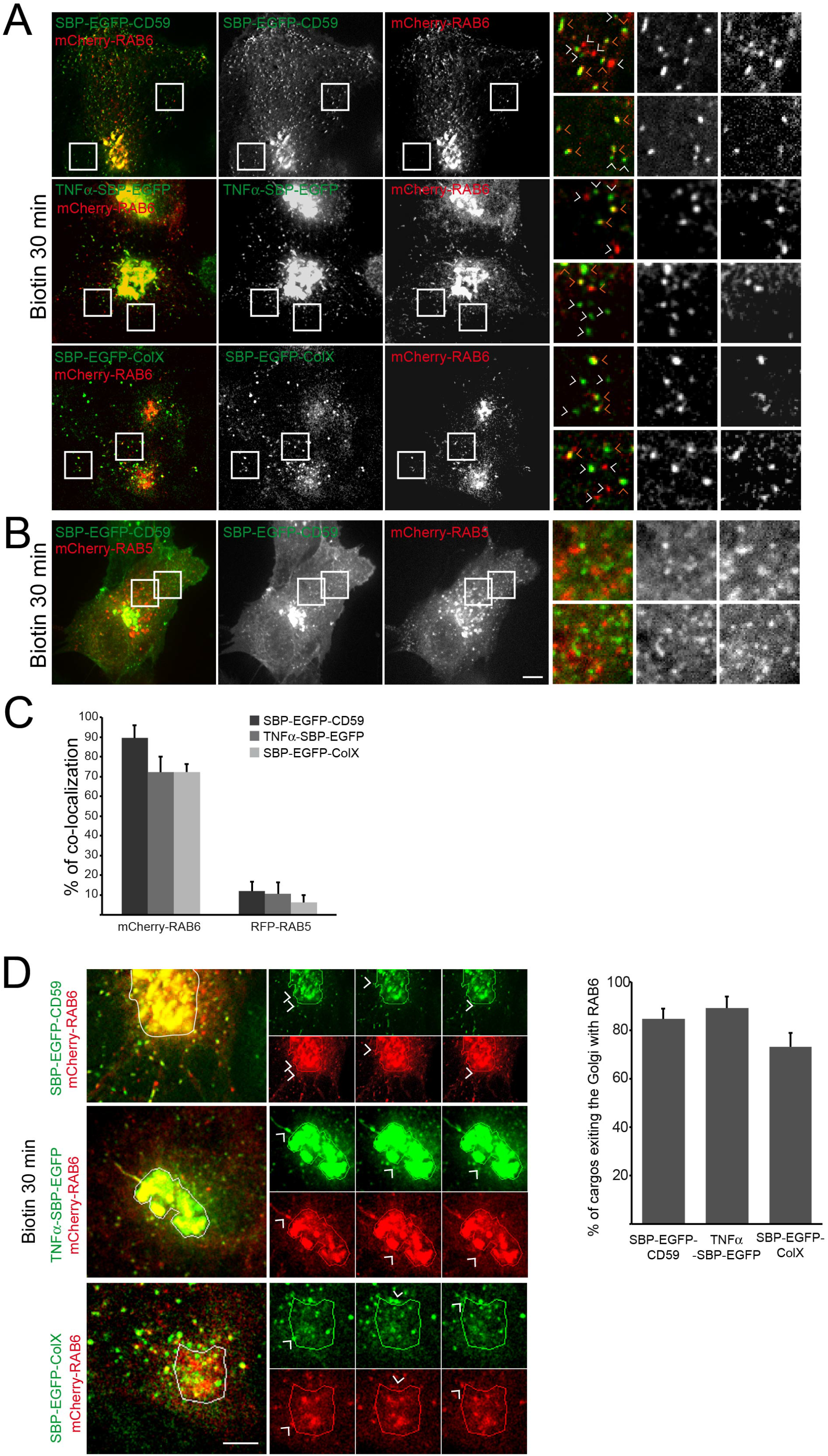
RAB6 associates with post-Golgi carriers containing CD59, TNFα or ColX. (**A**) RPE1-SBP-EGFP-CD59, HeLa-SBP-EGFP-ColX (stably expressing cells or transiently transfected cells), and HeLa-TNFα-SBP-EGFP cells co-expressing mCherry-RAB6 were incubated for 30 min with biotin to allow cargo release from the ER. Cells were imaged using a time-lapse spinning-disk confocal microscope. Representative images taken from movies are displayed. Orange arrowheads point at colocalized vesicles. White arrowheads point at not colocalized vesicles. (**B**) HeLa cells co-expressing SBP-EGFP-CD59 and RFP-RAB5A were incubated for 30 min with biotin to allow cargo release from the ER. Cells were imaged using a time-lapse spinning-disk confocal microscope. Representative images taken from movies are displayed. Scale Bar, 5 μM. (**C**) Quantification of the co-localization between mCherry-RAB6 or m-Cherry-RAB5 and each type of cargo (mean ± SEM, n= 9-24 cells). Scale Bar, 10 μM. (**D**) HeLa cells co-expressing mCherry-RAB6 and SBP-EGFP-CD59, TNFα-SBP-EGFP or SBP-EGFP-ColX were incubated for 30 min with biotin and imaged using time-lapse video-microscopy. Left panel: representative images of vesicles positive for SBP-EGFP-CD59, TNFα-SBP-EGFP or SBP-EGFP-ColX and mCherry-RAB6 exiting the Golgi complex together. Higher magnification of the images taken from time-lapse movies are on the right; Right panel: Quantification of the percentage of EGFP-SBP-CD59 positive vesicles exiting the Golgi complex with RAB6 (mean ± SEM, n= 13-18 cells). Scale bars, 10 μm.

To rule out the possibility that the high percentage of RAB6-positive post-Golgi transport carriers is due to RAB6 overexpression, the anti-RAB6:GTP antibody AA2 (Nizak et al., 2003) was used to analyze endogenous RAB6 expression. 60% co-localization was obtained between endogenous RAB6:GTP and either SBP-EGFP-CD59, TNFα-SBP-EGFP or SBP-EGFP-ColX present in post-Golgi vesicles (Supplementary Figure 3), thus indicating that the high percentage of post-Golgi transport carriers positive for RAB6 was not a consequence of RAB6 overexpression.

To investigate when RAB6 associates with post-Golgi carriers, we carefully examined them at the exit of the Golgi. As shown in Figure 4D, about 80% of vesicles containing SBP-EGFP-CD59, TNFα-SBP-EGFP or SBP-EGFP-CoIX exiting the Golgi complex were positive for RAB6. Importantly, this percentage of co-localization is similar to the 80% colocalization we found above when looking at the whole population of transport carriers.

Altogether, our results show that RAB6 is present on transport vesicles, irrespective of the transported cargo, when they leave the Golgi and remains associated with them until they reach the plasma membrane.

### The RAB6 machinery is required for the secretion of GPI-APs, TNFα and ColX

To address the functional role of RAB6, we assessed the effect of siRNA-mediated knockdown of RAB6 on the secretion of various cargos. A 50 % inhibition of the arrival of TNFα at the plasma membrane was observed 30 min and 60 min after biotin addition under conditions of RAB6 depletion as quantified using SPI (Figure 5A). Similarly, in the absence of RAB6, a 50 % reduction of ColX secretion was observed 120 min after biotin addition as analyzed by western-blotting of cell lysates and culture medium (Figure 5B). The arrival at the plasma membrane of placenta alkaline phosphatase (PLAP) was also delayed (Supplementary Figure 4A).

**Figure 5:**
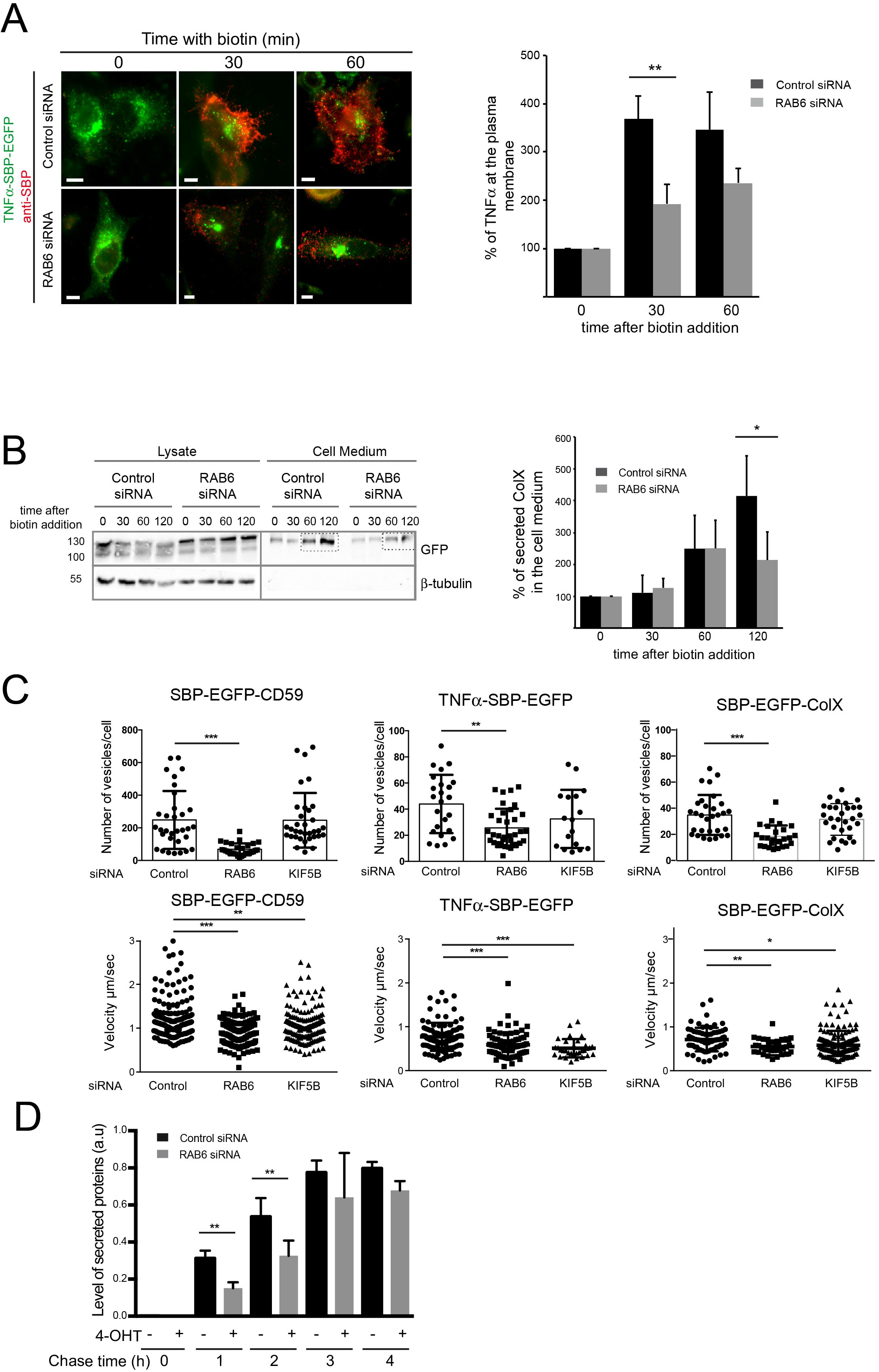
RAB6 and KIF5B are required for the proper secretion SBP-EGFP-CD59, TNFα-SBP-EGFP, SBP-EGFP-ColX at the plasma membrane. RAB6 depletion delays total protein secretion. (**A**) Left: HeLa-TNFα-SBP-EGFP, treated or not with RAB6 siRNAs, were incubated for 30 or 60 min with biotin to allow cargo release. The amount of secreted TNFα was determined on images acquired on cells using the SPI assay. Representative images of cells expressing TNFα-SBP-EGFP (green) and stained with anti-SBP (red) are displayed; Right: Quantification of the amount of TNFα present at the plasma membrane in cells treated as indicated above (mean ± SEM, n= 67-111 cells). ** p< 10^−2^ (Student’s t-test). Scale bars, 5 μm. (**B**) HeLa-SBP-EGFP-ColX treated or not with RAB6 siRNAs were incubated for 30, 60 or 120 min with biotin to allow cargo release from the endoplasmic reticulum. The amount of secreted cargo was revealed by western-blotting using an anti-GFP antibody. Beta-tubulin was used as a loading control and as a non-secreted protein. Representative immunoblots are displayed. Quantification of the amount of ColX present at the plasma membrane in cells treated as indicated above (mean ± SEM, n= 3). * p< 0.05 (Student’s t-test). (**C**) HeLa cells expressing SBP-EGFP-CD59, TNFα-SBP-EGFP and SBP-EGFP-ColX were treated for 3 days with control, RAB6 of KIF-5B siRNAs. Cells were incubated for 30 min with biotin to allow cargo release from the ER. Cells were imaged using a time-lapse spinning disk confocal microscope. Vesicles velocity (μm/sec) and the number of vesicles per cell were quantified using ImageJ (mean ± SEM, n= 17-35 cells). *** p< 10^−4^, ** p< 0,005, * p< 0.05 (Student’s t-test). (**D**) The SUnSET assay was used to determine the effect of RAB6 depletion on total protein secretion. Mouse embryonic fibroblasts (MEFs) prepared from RAB6 loxP/Ko Rosa26CreERT2-TG embryos (described in (Bardin et al., 2015)) (named after RAB6-KO) were treated with ethanol or with 4-hydroxytamoxifen (4-OHT) for 96 h to induce RAB6 depletion. Cells were then incubated with puromycin and chased in puromycin-free medium for 0, 1, 2, 4, 5.5 h. Total protein content in cell lysis or supernatant were labelled with an anti-puromycin antibody. Quantification of puromycin intensity in the supernatant and the whole cell lysis using Image Lab software (Bio-Rad). (mean ± SEM, n= 4). ** p< 0.05 (Student’s t-test). Representative immunoblot is displayed in Supplementary Figure 5.

RAB6 depletion leads to a 50% decrease in the number of post-Golgi vesicles containing VSV-G (Grigoriev et al., 2007; Miserey-Lenkei et al., 2010), which is similar to the reduction we observed in the number of SBP-EGFP-CD59, TNFα-SBP-EGFP or SBP-EGFP-ColX-positive post-Golgi vesicles (Figure 5C). This effect of RAB6 depletion results from a decreased recruitment on Golgi/TGN membranes of myosin II, whose activity is required for the fission of RAB6-positive vesicles from Golgi/TGN membranes (Miserey-Lenkei et al., 2010). To confirm the involvement of myosin II, cells expressing SBP-EGFP-CD59, TNFα-SBP-EGFP or SBP-EGFP-ColX were incubated with para-nitroblebbistatin for 15 min and then 30 min with biotin. As illustrated in Supplementary Figure 4B, SBP-EGFP-CD59 could be detected in membrane tubules connected to the Golgi that represent transport carriers that cannot detach from Golgi/TGN membranes (Miserey-Lenkei et al., 2010).

Finally, RAB6-positive vesicles are transported from the Golgi to the plasma membrane along microtubules by the KIF5B kinesin motor (Grigoriev et al., 2007; Miserey-Lenkei et al., 2010). Accordingly, cells treated with KIF5B siRNA showed a reduced velocity of CD59-, TNFα- or ColX-positive post-Golgi vesicles (Figure 5C).

Altogether the above results show that a variety of cargos use the RAB6 machinery for transport from Golgi to the plasma membrane. They also raise the possibility that RAB6 might be involved in the secretory process of all proteins leaving the Golgi complex. To test this hypothesis, we performed experiments on MEF cells derived from *RAB6* conditional KO mice embryos (Bardin et al., 2015) and global protein secretion was monitored using the SUnSET assay (Schmidt et al., 2009). MEFs either treated with 4-hydroxytamoxifen (4-OHT) to deplete RAB6 or with ethanol as a control (Supplementary Figure 5) were incubated with puromycin and then chased in puromycin-free medium for different time points. The amount of protein in cell lysates and supernatants was analyzed by western-blot using an anti-puromycin antibody and the relative amount of protein secreted quantified (Figure 5D; Supplementary Figure 5). To assess that the SUnSET assay worked properly, the same experiments were performed following incubation with the protein synthesis inhibitor cycloheximide, and as expected, no proteins were detected in both cell lysates and supernatants (Supplementary Figure 5). Following RAB6 depletion, the total amount of secreted protein after a 1 h or 2 h chase was reduced by 50% although total protein expression was unaffected (see cell extracts at t=0 of cell treated or not with 4-OHT, Supplementary Figure 5). As apparent on the western blots, the intensity of all secreted protein bands was decreased upon RAB6 depletion, suggesting a global effect on protein secretion (Figure 5D). Importantly, after 4 h and 5.5 h of chase, the amount of protein secreted by cells expressing or not RAB6 was similar (Figure 5D). These results thus showed that RAB6 depletion does not block the secretory process but leads to a delay in total protein secretion, as previously found for exogenous cargos (Grigoriev et al., 2007; Miserey-Lenkei et al., 2010).

Altogether, the above results show that RAB6 function is required for total protein secretion at the cell level.

### RAB6 is not involved in sorting of cargos at the exit of Golgi complex

Newly synthesized proteins are thought to be sorted into distinct populations of transport carriers at the TGN (De Matteis and Luini, 2008). The RUSH system allows to synchronize transport of two (or more) cargos at the same time, thus providing a powerful approach to investigate sorting processes in further detail. We imaged cells expressing two cargos, either SBP-EGFP-ColX and TNFα-SBP-mCherry, SBP-EGFP-ColX and SBP-mCherry-CD59, or TNFα-SBP-EGFP and SBP-mCherry-CD59 (Figure 6A). In all cases, although a majority (60%) of the vesicles contained two cargos, a large fraction of vesicles contained only one cargo. This indicates that, despite the sudden wave of transport imposed by biotin-induced release from the ER, efficient sorting can still occur at the Golgi complex. We then estimated the percentage of co-localization between RAB6 and post-Golgi vesicles containing one or two cargos. As illustrated in Figure 6B in the case of TNFα and ColX, the majority (60%) of vesicles containing the two cargos was positive for endogenous RAB6. Importantly, similar percentages of vesicles containing only one cargo were positive for RAB6 (50% and 60% of vesicles containing ColX or TNFα, respectively) (Figure 6B). This suggests that RAB6 is not involved in sorting of cargos at the exit of the Golgi complex but associates to transport carriers, irrespective of the transported cargo, to target them to specific sites at the plasma membrane.

**Figure 6:**
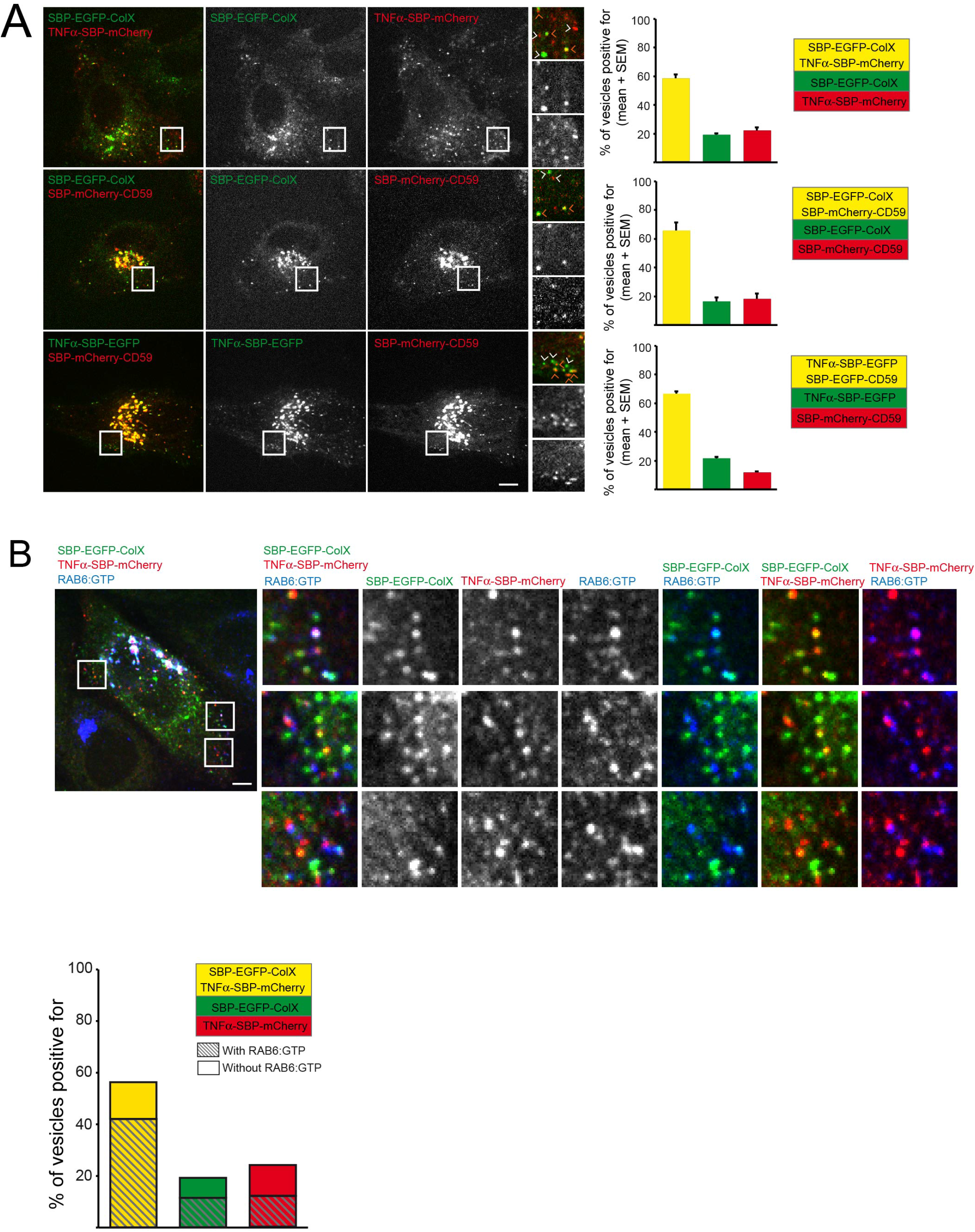
RAB6 is not involved in sorting of cargos at the exit of Golgi complex. (**A**) HeLa cells co-expressing SBP-EGFP-ColX and TNFα-SPB-mCherry, SBP-EGFP-ColX and SBP-mCherry-CD59 or TNFα-SBP-EGFP and SBP-mCherry-CD59 were incubated for 35 min with biotin to allow cargo release from the ER and then imaged using time-lapse video-microscopy. Representative images taken from movies are shown (higher magnification on the right). Orange arrowheads point at colocalized vesicles. White arrowheads point at not colocalized vesicles. Right panel: Quantification for each pair of cargos of the percentage of vesicles positive both cargos (yellow), EGFP (green) or mCherry (red) (mean ± SEM, n= 197-501 vesicles from 11 to 20 cells). (**B**) Staining of endogenous RAB6:GTP (blue) in cells co-expressing SBP-EGFP-ColX (green) and TNFα-SBP-mCherry (red) and incubated for 30 min with biotin to visualize post-Golgi carriers. Higher magnifications are shown on the right. Percentage of vesicles containing both SBP-EGFP-ColX and TNFα-SPB-mCherry (yellow), only SBP-EGFP-ColX (green) or only TNFα-SPB-mCherry (red). Dashed bars indicate the percentage of vesicles positive for RAB6. (mean ± SEM, n= 24-128 vesicles). Scale bars: 5 μm.

## DISCUSSION

Cells need to continuously control the transport of various proteins to the cell surface both to ensure homeostasis and to sustain differentiated functions. The study of various transport steps, like ER to Golgi or Golgi to ER, have shown that mechanisms are at work which enable the transport of a diversity of proteins using a “universal” core machinery (COPI, COPII for example). However, whether Golgi to plasma membrane is similarly controlled by a core machinery was still unclear. One of the main findings of this study is that post-Golgi transport of diverse secretory cargos, of various shape, function and characterized by diverse transport kinetics are handled by a common machinery and reach the membrane in similarly restricted domains.

The concept of “exocytosis hotspots” existing in cells was actually proposed almost 40 years ago by D. Louvard (Louvard, 1980) when analyzing the secretion of fibronectin after cell attachment (Heggeness et al., 1978). Surprisingly, we found that the secretion of cargos recently released from the Golgi complex occurs in so called “hotspots” close to focal adhesions, and that this is a general process common to several and diverse proteins.

Targeting to focal adhesions does not seem to depend on the particular function or modification of the cargo because it was observed for glycosylated (gp135 and Collagen X) and non-glycosylated proteins (TNFα). This was also observed for soluble secretory GFP, which is non-glycosylated and unlikely to bear any specific transport signal. The secretion of particular cargo types, like metalloproteinases such as MT1-MMP, at adhesion sites was previously proposed (Stehbens et al., 2014; Bravo-Cordero et al., 2007). However, it was not clear if the vesicles containing MT1-MMP were strictly secretory vesicles or vesicles recycled from the endocytic/recycling pathways. Here, by synchronizing the anterograde transport of cargos using the RUSH assay, we unambiguously analyzed the first wave of secretion of transport carriers. The SPI assay that we set-up for capture of secreted proteins prevented transmembrane diffusion in the plasma membrane or release of soluble cargos into the extracellular space. The SPI provides insight into the secretion history of synchronized cargos. By combining these two assays, we were able to show that transport carriers use preferential microtubule tracks to reach secretion hotspots. In addition to this directed movement, the exocytic zones may be restricted close to focal adhesions by the presence of factors necessary for docking/fusion of transport carriers such as ELKS and LL5β proteins (Grigoriev et al., 2011).

Focal adhesions both produce and sense adhesion forces. They are able to modulate cell adhesion in response to cellular environment, for instance by modifying attachment to the extracellular matrix. At the intracellular level, the interaction between microtubules and focal adhesions impacts the structure and dynamics of focal adhesions. This interaction is essential for cellular organization, polarization and migration (see (van der Vaart et al., 2009) for review). Here we observed that their localization is correlated to the existence of secretion hotspots. Since we could not observe the early steps of focal adhesion establishment while monitoring exocytosis, it is not possible to precisely decipher the mechanisms allowing focal adhesions and exocytosis to occur in these domains. Focal adhesions could dictate the presence of secretion hotspots, restricted exocytosis domains could favor focal adhesions, or upstream mechanisms such as local absence of cortical actin or the accumulation of adhesion proteins in plasma membrane sub-domains could enable focal adhesions and exocytosis to occur in these hotspots.

An important finding of this study is that RAB6 is likely a major regulator of post-Golgi secretion (Figure 7). RAB6 was previously shown to be present on post-Golgi transport carriers containing VSV-G, NPY and TNFα (Grigoriev et al., 2007; Miserey-Lenkei et al., 2010; Micaroni et al., 2013; Johns et al., 2014). Here we show that RAB6 associates with secretory vesicles containing a variety of exogenous cargos and that its depletion delays their arrival at the plasma membrane. The observation that the secretion of most endogenous proteins synthesized by MEF cells is affected by RAB6 depletion further suggests that RAB6 may be present on the majority of post-Golgi vesicles transporting cargos to the plasma membrane. This is in agreement with a model proposing that one of the main functions of RAB6 is to target post-Golgi vesicles to cortical ELKS-containing patches that define secretion domains (Grigoriev et al., 2007; 2011). ELKS may not only dock RAB6-positive vesicles to the plasma membrane but also to other compartments. We have recently shown that ELKS is present on the membrane of mature melanosomes in melanocytes, which allows the diversion of part of the RAB6-dependent secretory pathway to melanosomes, enabling direct transport of a subset of melanosomal enzymes from Golgi (Patwardhan et al., 2017).

**Figure 7:**
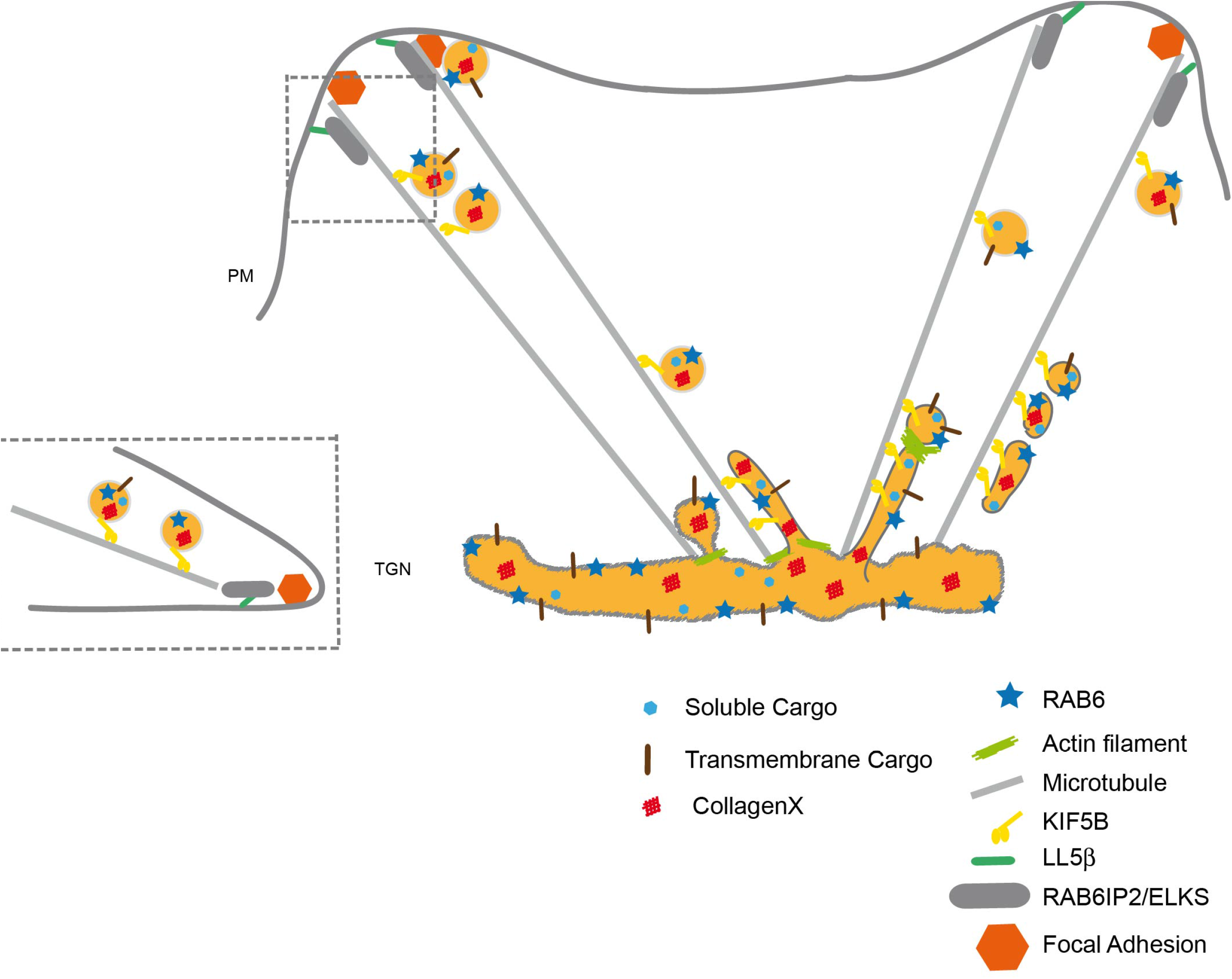
Proposed model to explain the role of RAB6 in post-Golgi secretory pathway. All RAB6 effectors associated to post-Golgi secretion, myosin II, KIF5B and ELKS/RAB6IP2, are involved in the secretion of various cargos. The different events can be envisioned as follows. RAB6 is present on CD59-, TNFα- and ColX-containing vesicles exiting the Golgi complex. Myosin II is implicated in the fission of these vesicles from Golgi membranes (Miserey-Lenkei et al., 2010). Vesicles are transported to the plasma membrane in a KIF5B-dependent manner (Grigoriev et al., 2007; Miserey-Lenkei et al., 2010) and ELKS ensures their docking to exocytic hotspots close to focal adhesions (Grigoriev et al., 2007; 2011).

It is likely that RAB6 plays another role beyond its function in targeting post-Golgi vesicles to secretion hotspots. We have previously shown that RAB6 not only controls the fission of its own transport carriers (Miserey-Lenkei et al., 2010) but also could select a subset of MTs originating from Golgi membranes for their transport (Miserey-Lenkei et al., 2017). Although not directly proven, it is likely that this subset of MTs correspond to the ones identified in this study which are used several times for targeting cargos to secretion hotspots.

Evidence exist that several classes of secretory carriers can form at the trans-Golgi, indicating that cargos are sorted prior to be delivered to the plasma membrane (De Matteis and Luini, 2008). Our study highlights that, even when pulsing a massive amount of cargos in the secretory pathway, the Golgi is still able to properly sort a large percent of cargos in dedicated transport intermediates. Such sorting may be even more important in polarized cells, such as epithelial or neuronal cells, or when studying cargos targeted to particular intra-cellular compartments like the endo-lysosomal system. Such an intra-Golgi sorting is also evident when monitoring the time of residence in the Golgi and exit from the Golgi of cargo. Some cargos exit very quickly while others stay for a long time in the Golgi complex, suggesting intra-Golgi segregation. However, our data indicate that irrespective of the cargo, the vast majority of transport intermediates are positive for RAB6. This suggests that RAB6 is not involved in cargo sorting but belongs to a general machinery responsible for fission of transport intermediates from the Golgi (Miserey-Lenkei et al., 2010) and their fusion with target compartments.

In conclusion, our study highlights the existence of secretion hotspots that occur near focal adhesions, and indicates RAB6 as a central actor in these mechanisms. Why localized exocytosis domains exist remains an open question and will require further work. These domains may simply reflect the global architecture of the cell dictated by the organization of the actin and microtubule networks. Alternatively, they may result from functional links existing between Golgi membranes and focal adhesions, which are important for cell adhesion, polarization and migration.

## MATERIAL AND METHODS

### Cells

HeLa and RPE-1 cells were grown at 37°C with 5% CO_2_ in Dulbecco’s Modified Eagle Medium (DMEM, high glucose, GlutaMAX, Life Technologies) or DMEM-F12 (Gibco), respectively, supplemented with 10% fetal calf serum (FCS, GE Healthcare and Eurobio), pyruvate sodium 1mM (Life Technologies), penicillin and streptomycin 100 U/mL (Life technologies and Gibco). HeLa cells stably expressing RUSH constructs were cultured as described previously (Fourriere et al., 2016) in medium supplemented with 4 μg/ml of puromycin (Invitrogen). Lentiviral infection was used to generate RPE-1 cells stably expressing Streptavidin-KDEL and SBP-EGFP-CD59. Mouse embryonic fibroblasts (MEFs) were prepared from mouse RAB6 loxP/loxP (control) or RAB6 loxP/KO Rosa26CreERT2-TG (RAB6 KO) mouse embryos (Bardin et al., 2015). MEFs cells were grown in DMEM (Gibco) supplemented with 10% fetal calf serum (FCS) (Eurobio) and 100 U/mL penicillin/streptomycin (Gibco). For RAB6 depletion, MEFs cells were incubated with 1 μM 4-hydroxytamoxifen (4-OHT) for 96 h (Sigma).

### Plasmids

The plasmids encoding the following fusion proteins were used: GFP-/mCherry RAB6A or -RAB6A’ (Miserey-Lenkei et al., 2010); GFP-ELKS (a gift from A. Akhmanova, Utrecht University, The Netherlands); VSV-GtsO45-EGFP (Hirschberg et al., 1998). Paxillin-BFP, paxillin-GFP and paxillin-mCherry were constructed from the Human paxillin cDNA from Open Biosystems (Accession number BC144410). All the RUSH plasmids used in this study (except VSV-G-SBP-EGFP) use Streptavidin-KDEL as a hook. VSV-G-SBP-EGFP uses Streptavidin-Ii as a hook. Briefly, the hook (Streptavidin-tagged protein) allows anchoring of the SBP-tagged reporter in the endoplasmic reticulum in the absence of biotin thanks to Streptavidin-SBP interaction. RUSH plasmids coding for TNFα-SBP-EGFP, TNFα-SBP-mCherry, TNFα-SBP-BFP, SBP-EGFP-E-cadherin, SBP-ssEGFP were previously described (Boncompain et al., 2012; Fourriere et al., 2016). The other RUSH plasmids used were generated as previously described (Boncompain and Perez, 2013). The accession numbers of the corresponding reporter used as template are the followings: CollagenX (BC130621), gp135/podocalyxin (NP_001076235), CD59 (NP_000602), PLAP (NP_001623). The release of the RUSH cargoes was induced by addition of 40 μM D-biotin (Sigma).

### Biochemical reagents

The following reagents were used: Para-nitroblebbistatin (Optopharma Ltd), 4-OH tamoxifen (Sigma). SiR-tubulin (Spirochrome) was used to label microtubule in living cells according to manufacturer’s instructions.

### DNA and RNA transfection

HeLa or RPE-1 cells were transfected 24 to 48 h before observation with calcium phosphate (Jordan et al., 1996) or with X-tremeGEN9 (Roche), following the manufacturer’s instructions. For RNA interference experiments, cells were transfected with the corresponding siRNA (RAB6A/A’, KIF5B, ELKS or Luciferase) using Lipofectamine RNAiMAX (Invitrogen), following the manufacturer’s instructions. The sequences of the siRNAs used in this study are the following: siRNA Luciferase (5’-CGUACGCGGAAUACUUCGA-3’, Sigma); siRNA against Human RAB6 targeting RAB6A/A’ (Del Nery et al., 2006) (5’-GACAUC UUU GAU CAC CAGA-3’, Sigma); siRNA against Human KIF5B (Grigoriev et al., 2007) (5’-AACGTTGCAAGCAGTTAGAAA-3’, Sigma); siRNA against ELKS (Grigoriev et al., 2007) (5’-GUGGGAAA ACCCUUUCAAU-3’, Ambion).

### Specific protein immobilization assay (SPI)

Coverslips were incubated in sterile conditions in a bicarbonate solution (0,1 M, pH 9,5, 1 h, 37°C) followed by incubation in poly-L lysine 0.01% (diluted in water, 1 h, 37°C, Sigma). Coverslips were then washed in PBS and dried before antibody incubation (diluted in the bicarbonate buffer) 3 h at 37°C (or overnight). Coverslips were washed twice in PBS and then with cell medium before cells were seeded. Antibodies used for coating in this study were: anti-GFP (A-P-R#06 from the Recombinant Antibody Platform of the Institut Curie, dilution 1:400), anti-mCherry (A-P-R#13 from Recombinant Antibody Platform, dilution 1:400), anti-VSV-G (a gift from T. Kreis, University of Geneva, dilution 1:50) and anti-SBP (Millipore, MAB10764, batch 2697549, dilution 1:60).

### Immunofluorescence

Cell fixation was performed with paraformaldehyde 3% or 4% (Electron Microscopy Sciences) for 15 min at room temperature. For permeabilization, cells were incubated in PBS supplemented with BSA 2 g/l and saponin 0.5 g/l for 10 min at room temperature. Surface staining was performed at 4°C on non-fixed intact cells incubated with the primary antibody for 40 min. Cells were then fixed with paraformaldehyde 2% for 10 min at room temperature. Primary antibodies used in this study were mouse anti-GFP (Roche catalog number 11814460001, batch 11063100, dilution 1:1000), rabbit anti-paxillin (Abcam ab32084 batch GR23669-20, dilution 1:250), mouse anti-VSV-G (a gift from T. Kreis, University of Geneva, dilution 1:500), mouse anti-GM130 (BD Biosciences, catalog number 610823, batch 4324839, dilution 1:1000), Human anti-RAB6:GTP (AA2, Adipogen, dilution 1:250); mouse monoclonal anti-SBP tag (clone 20, Millipore, dilution 1:400); rabbit polyclonal anti-GFP (recombinant antibody platform of the Institut Curie, dilution 1:400);; monoclonal mouse anti-GFP (Roche, dilution 1:1000); Human anti-α-tubulin (Sigma, dilution 1:1000), polyclonal rabbit anti-RAB6 (Santa Cruz, dilution 1:1000); rabbit anti-ELKS (1:1000) (Monier et al., 2002), ubiquitous kinesin heavy chain anti-KIF5B (Santa Cruz Biotechnology, 1:1000). Alexa- and HRP- coupled secondary antibodies were purchased from Jackson ImmunoResearch Laboratory. Coverslips were mounted in Mowiol and examined under a 3D deconvolution microscope (Leica DM-RXA2), equipped with a piezo z-drive (Physik Instrument) and a 100×1.4NA-PL-APO objective lens for optical sectioning. 3D or 1D multicolor image stacks were acquired using the Metamorph software (MDS) through a cooled CCD-camera (Photometrics Coolsnap HQ).

### Live cell imaging

Spinning-disk confocal time-lapse imaging were done at 37°C in a thermostat-controlled chamber using an Eclipse 80i microscope (Nikon) equipped with spinning disk confocal head (Perkin), a 100x objective and either a Ultra897 iXon camera (Andor) or CoolSnapHQ2 camera (Roper Scientific). Fixed sample were on the other hand imaged using a 60x objective using the same setup. TIRF microscopy was done using Leibovitz’s medium (Life Technologies) at 37°C in a thermostat-controlled chamber. An Eclipse Ti inverted microscopes (Nikon) equipped with either a TIRF module (Nikon) or an iLAS2 azimuthal TIRF module (Roper Scientific), a 100x TIRF objective, a beam splitter (Roper Scientific) and an Evolve 512 EMCCD camera (Photometrics) was used in this case (Boulanger et al., 2014). All acquisitions were driven by Metamorph (Molecular Devices).

### Fluorescent activated cell sorting (FACS)

Cells were incubated at 4°C with an anti-GFP antibody diluted in PBS (Recombinant protein platform, Institut Curie, 1:400). Intra- and extra-cellular EGFP intensity were detected with a BD Accuri™ C6 Cytometer. The ratio between the cell surface signal intensity per the total signal intensity detected in EGFP-PLAP expressing cells measured at 30 and 60 minutes was normalized to the cell surface signal intensity detected at 0 minute (corresponding to background) for each condition.

### SUnSET assay (SUrface SEnsing of Translation)

SUnSET assay was carried out as described previously (Schmidt et al., 2009). Briefly, MEF cells were seeded at a confluence of 30 % then treated with ethanol (control) or 1 μM of 4-OH Tamoxifen for 96 h to deplete RAB6. The day of the experiment, cells were cultured without serum and incubated for 30 min at 37 °C with 10 μg/ml of puromycin. Puromycin was then chased at different time points (0, 1, 2, 4, and 5.5 h). The supernatants and the whole cell lysates were collected and processed for western-blotting. The puromycin signal was revealed using an anti-puromycin antibody (clone 12D10 Millipore MABE343). Quantification of puromycin intensity in the supernatant and the whole cell lysis from four independent experiment using Image Lab software (Bio-Rad) is shown. The amount of total secreted proteins was determined by the normalization of the intensity of puromycin signal in the supernatant for each time point by the sum of the intensity of puromycin signal in the supernatant and the whole cell lysate for each corresponding time point.

### Cell lysis, SDS-PAGE, and Western-blotting

Cells were washed three times in ice-cold PBS, scraped and then lysed in a buffer containing 150 mM NaCl, 50 mM Tris-HCl pH 7.5, 1% Nonidet-P40 (Sigma). Protein concentrations were determined by Quick Start™ Bradford 1x Dye Reagent (Bio-Rad). Equal amounts of proteins were reduced with 1x loading buffer containing 6% β-mercaptoethanol and resolved on 10% SDS-PAGE. Proteins were transferred onto nitrocellulose Protran BA 83 membrane (Life science), processing for immunoblotting. HRP-conjugated secondary antibodies associated signal was detected with ECL system and an enhanced chemiluminescence system (ChemiDoc Touch System, Bio-Rad). Quantification of the corresponding signal obtained was done using Image lab software.

ColX secretion following RAB6 depletion was measured using western-blotting. After release of the cargo from the ER for different time points, trafficking was stopped by incubating the cells at 4°C and the supernatant was collected. Secreted collagen X was concentrated using VIVASPIN TURBO 4 (filters <10kDa) column. When the culture media volume reached 65 μl, the centrifugation was stopped and 5X-loading buffer containing 6% β-mercaptoethanol was added. Cells were lysed in 150 mM NaCl, 50 mM Tris-HCl pH 7.5, 1 % Nonidet-P40 (Sigma) and proteins were boiled at 95 °C for 5 min. Samples were then processed for western-blotting.

### Image analysis and quantification

For Figures 1A, 2D, 2E and 2F, temporal projections were performed with the tool “Temporal-color code” on the Fiji software. For Figures 2G-H intensity profiles were calculated from a drawing line with the Fiji software. The proximity map of Figure 2C was done by Jérôme Boulanger. Distance from the paxillin marker (x) was calculated at each position in function of the frequency of the secretion puffs (y). Kymographs of the Figures 2E, 2G and 2H were done on the Fiji software. Quantification of co-localization in post-Golgi carriers (Figures 4C, 4D, 6A, 6B, S2, S3) was performed manually. Colocalization between two moving vesicles (or fixed vesicles in Figure 6B) in two different channels was determined manually using Image J software (Synchronize windows). In Figure S3B, colocalization was measured with Pearson’s coefficient using Image J software. Quantification of the velocity and the track length (the covered distance) of vesicles were done with the plug-in “Manual Tracking” from ImageJ software developed by Fabrice Cordelières (Institut Curie). The number of vesicles was determined using Image J software, using the “find maxima” plug-in. The number of vesicles was normalized either per cell or per area in each cell.

### Statistical analysis

All data were generated from cells pooled from at least 3 independent experiments represented as (*n*), unless mentioned, in corresponding legends. Statistical data were presented as means ± standard error of the mean (S.E.M.). Statistical significance was determined by Student’s t-test for two or three sets of data using Excel, no sample was excluded. Cells were randomly selected. Only P-value <0.05 was considered as statistically significant.

## ACKNOWLEDGEMENTS

The authors acknowledge Shauna Katz for careful reading of the manuscript. The authors gratefully acknowledge the Cell and Tissue Imaging Facility (PICT-IBiSA), Institut Curie, a member of the French National Research Infrastructure, France-BioImaging (ANR10-INBS-04). The authors also thank the recombinant antibody platform of the Institut Curie. Work performed in F.P. laboratory was funded by CNRS (Centre National pour la Recherche Scientifique), the Fondation pour la Recherche médicale (FRM DEQ20120323723 and FRM FDT20150532154 to L.F.), the Labex CellTisPhyBio (L.F.), and the Agence Nationale de la Recherche (ANR-12-BSV2-0003-01). Work performed in B.G. laboratory was supported by an ERC (European Research Council) advanced grant (project 339847 ‘MYODYN’), ANR (To A.K.), ARC (to A.K.) and Institut Curie (to A.K.). The Goud and Perez teams are member of Labex CelTisPhyBio (11-LBX-0038) and Idex Paris Sciences et Lettres (ANR-10-IDEX-0001-02 PSL).

## Authors Contributions

L.F. and N.G. carried out the experiments presented in Figs. 1, 2, 3a-c, Supplementary Figure. 1; A.K. the experiments in Fig. 3d-g, 4, 5, 6 and Supplementary Figs. 2-5; S.B. the experiments in Supplementary Fig 5; J.B and R.S. helped for image analysis and TIRF; G.B. designed and generated the RUSH plasmids; L.F., A.K., S.B., N.G., G.B., S.M-L., F.P., B.G. designed and interpreted the experiments; L.F., A.K., G.B., S.M-L., F.P., B.G. wrote the manuscript. L.F., A.K., S.B., N.G., J.B., R.S., G.B., S.M-L., F.P., B.G. edited the manuscript. B.G and F.P. secured funding.

